# Analysis of centrosomal area actin reorganization and centrosome polarization upon lymphocyte activation at the immunological synapse

**DOI:** 10.1101/2021.09.29.462395

**Authors:** Sara Fernández-Hermira, Irene Sanz-Fernández, Marta Botas, Víctor Calvo, Manuel Izquierdo

## Abstract

T cell receptor (TCR) and B cell receptor (BCR) stimulation of T and B lymphocytes, by antigen presented on an antigen-presenting cell (APC) induces the formation of the immunological synapse (IS). IS formation is associated with an initial increase in cortical filamentous actin (F-actin) at the IS, followed by a decrease in F-actin density at the central region of the IS, which contains the secretory domain. This is followed by the convergence of secretion vesicles towards the centrosome, and the polarization of the centrosome to the IS. These reversible, cortical actin cytoskeleton reorganization processes occur during lytic granule secretion in cytotoxic T lymphocytes (CTL) and natural killer (NK) cells, proteolytic granules secretion in B lymphocytes and during cytokine-containing vesicle secretion in T-helper (Th) lymphocytes. In addition, several findings obtained in T and B lymphocytes forming IS show that actin cytoskeleton reorganization also occurs at the centrosomal area. F-actin reduction at the centrosomal area appears to be associated with centrosome polarization. In this chapter we deal with the analysis of centrosomal area F-actin reorganization, as well as the centrosome polarization analysis towards the IS.

## 1 Introduction

### 1.1 The immunological synapse

T and B lymphocyte activation by APC takes place at a specialized cell to cell interface called the IS. APC have the ability to present antigens to T lymphocytes bound to major histocompatibility complex (MHC) molecules (Huppa and Davis, 2003) (Fooksman et al., 2010). In contrast, B lymphocytes can directly recognize antigen tethered to the cell surface of specialized APC (Yuseff et al., 2013). In addition, NK cells were first noticed for their ability to kill tumor cells after IS formation, but without any priming or prior activation, in contrast to CTL, which need priming and activation by APC (Lagrue et al., 2013). IS establishment by T and B lymphocytes and NK cells is a very dynamic, plastic and critical event, acting as a tunable signalling platform that integrates spatial, mechanical and biochemical signals, involved in specific, cellular and humoral immune responses (Fooksman et al., 2010) (de la Roche et al., 2016). The general architecture of the IS is described by the formation of a concentric, bullseye spatial pattern, named the supramolecular activation complex (SMAC), upon cortical actin reorganization (Griffiths et al., 2010) (Billadeau et al., 2007) (Kuokkanen et al., 2015) (Yuseff et al., 2013) (Carisey et al., 2018). In the IS made by T and B lymphocytes, this reorganization yields a central cluster of antigen receptors bound to antigen called central SMAC (cSMAC) produced by centripetal traffic, and a surrounding adhesion molecule-rich ring, called peripheral SMAC (pSMAC), which appears to be crucial for adhesion with the APC (Monks et al., 1998) (Fooksman et al., 2010). The distal SMAC (dSMAC) is located surrounding the T and B lymphocytes pSMAC, at the edge of the contact area with the APC, and comprises a circular array of dense F-actin (Griffiths et al., 2010) (Le Floc’h and Huse, 2015) (Ritter et al., 2013) (Rak et al., 2011). More recently, several super-resolution imaging techniques have revealed that, upon TCR-antigen interaction, at least four discrete F-actin networks form and maintain the shape and function of this canonical IS (Hammer et al., 2018) (Blumenthal and Burkhardt, 2020).

### 1.2 Polarization of centrosome and secretory granules

IS formation induces the convergence of T and B lymphocytes and NK cells secretion vesicles towards the centrosome and, simultaneously, the polarization of the centrosome (the major microtubule-organizing center –MTOC– in lymphocytes) towards the IS (de la Roche et al., 2016) (Huse, 2012). These traffic events, acting together, lead to polarized secretion of extracellular vesicles and exosomes coming from multivesicular bodies (MVB) in T and B lymphocytes and NK cells (Peters et al., 1991) (Alonso et al., 2011) (Herranz et al., 2019) (Calvo and Izquierdo, 2020); lytic granules in CTL (Stinchcombe et al., 2006) (de la Roche et al., 2016); stimulatory cytokines in Th cells (Huse et al., 2008); lytic proteases in B lymphocytes (Yuseff et al., 2013).

### 1.3. Synaptic actin cytoskeleton regulation of secretory traffic

#### 1.3.1 Cortical actin cytoskeleton

Actin cytoskeleton reorganization plays a central role in IS maintenance, but also in antigen receptor-derived signalling in T and B lymphocytes (Billadeau et al., 2007) and activating receptor signalling in NK cells (Ben-Shmuel et al., 2021). Please refer to the excellent reviews on this subject in the IS made by B lymphocytes (Yuseff et al., 2009) (Yuseff et al., 2013), T lymphocytes (Billadeau et al., 2007) (Ritter et al., 2013), and NK cells (Lagrue et al., 2013) (Ben-Shmuel et al., 2021). The concentric F-actin architecture of the IS and the actin cortical cytoskeleton reorganization are shared by B lymphocytes, CD4^+^ Th lymphocytes, CD8^+^ CTL, and NK cells (Le Floc’h and Huse, 2015) (Brown et al., 2011). Remarkably, all these immune cells exhibit the ability to form synapses and to directionally secrete proteases, cytokines or cytotoxic factors at the IS. Thus, this polarized secretion in the context of the F-actin synaptic architecture, most probably, enhances the specificity and the efficacy of the subsequent responses to these factors (Le Floc’h and Huse, 2015) by spatially and temporally focusing the secretion at the synaptic cleft (Billadeau et al., 2007), which avoids the stimulation or death of bystander cells.

At the early phases of IS formation, F-actin accumulates at the contact area (called IS interface) of the immune cell with the APC, to generate filopodia and lamellipodia, that produce dynamic changes between extension and contraction in the lymphocyte over the surface of the APC (Le Floc’h and Huse, 2015). Subsequently, once IS evolution has stabilized, F-actin reduction from the cSMAC appears to facilitate secretion toward the APC by focusing secretion vesicles on the IS (Stinchcombe et al., 2006) and, almost simultaneously, cortical F-actin accumulates into the dSMAC. Thus, F-actin forms a permissive network at the IS of CTL and NK cells (Ritter et al., 2015) (Carisey et al., 2018). These reversible, cortical actin cytoskeleton reorganization processes occur during lytic granules secretion by both CTL and NK cells, but also during the polarization of some cytokine-containing secretion vesicles in Th lymphocytes (Griffiths et al., 2010;Chemin et al., 2012;Ritter et al., 2015), despite both the nature and cargo of the secretion vesicles in these cell types being quite different. Moreover, it has been shown that cortical F-actin density recovery at the IS leads to termination of lytic granule secretion in CTL (Ritter et al., 2017), supporting the role of actin cytoskeleton in initiation, but also in termination of granule secretion.

#### 1.3.2 Centrosomal actin cytoskeleton regulation of secretory traffic

F-actin reduction at the cSMAC does not simply allow secretion, as it apparently plays an active role in the initiation of centrosome and secretory granules movement towards the IS (Stinchcombe et al., 2006) (Ritter et al., 2015). In this context, some results suggest that cortical actin reorganization at the IS is necessary and sufficient for centrosome and lytic or cytokine-containing granules polarization (Ritter et al., 2015) (Chemin et al., 2012) (Sanchez et al., 2019).

However, other results show that the elimination of certain F-actin regulators such as formins FMNL1 or Dia1 inhibits centrosome polarization without affecting Arp2/3-dependent cortical actin reorganization (Gomez et al., 2007) supporting that, at least in the absence of FMNL1 or Dia1, cortical actin reorganization is not sufficient for centrosome polarization. Conversely, in the absence of cortical actin reorganization at the IS occurring in Jurkat T lymphocytes lacking Arp2/3, the centrosome can polarize normally to the IS (Gomez et al., 2007) (Kumari et al., 2014). All these results support that centrosome and secretory granules polarization induced by IS formation are regulated by HS1/WASp/Arp2/3-dependent cortical and formin-dependent non-cortical actin networks (Gomez et al., 2007) (Kumari et al., 2014). Thus, analysis of all these F-actin networks at different subcellular locations is necessary to achieve the full picture of the cellular actin reorganization processes leading to polarized secretion. Included among non-cortical actin networks, centrosomal area F-actin depletion appears to be crucial to allow centrosome polarization towards the IS in B lymphocytes stimulated with BCR-ligand-coated beads, used as a B lymphocyte synapse model (Obino et al., 2016). These data do not directly allow inferring either the sufficiency or the relative contribution of each F-actin network (cortical and centrosomal area) to centrosome polarization, unless specific experimental approaches are designed to address this point (Bello-Gamboa et al., 2020). The above-mentioned results on B lymphocytes, together with our results showing that PKCδ interference affects cortical F-actin at the IS (Herranz et al., 2019), prompted us to study the involvement of centrosomal area F-actin in centrosome polarization in cell-to-cell synapses made by T lymphocytes and its potential regulation by PKCδ (Bello-Gamboa et al., 2020). **The experimental approach described here allows the simultaneous evaluation of centrosome polarization and centrosomal area F-actin, at the single cell level**. To develop these analyses, modifications of the methods previously described in B lymphocytes to measure normalized centrosome polarization (Obino et al., 2017) (Saez et al., 2019) and centrosomal area F-actin (Obino et al., 2016) (Ibanez-Vega et al., 2019) were used. Following this approach, we have shown that, upon Th lymphocyte IS formation, centrosomal area F-actin decreased concomitantly with centrosome polarization to the IS, and a linear correlation between these two parameters exists (Bello-Gamboa et al., 2020).

## 2. Materials, cells, immunological synapse formation and image capture

1. Raji B and Jurkat T (clone JE6.1) cell lines are obtained from the ATCC.
2. Cell lines are cultured in RPMI 1640 medium containing L-glutamine (Invitrogen) with 10% heat-inactivated fetal calf serum (FCS) (Gibco) and penicillin/streptomycin (Gibco) and 10 mM HEPES (Lonza).
3. For IS formation, Raji cells are attached to Ibidi 8 microwell culture dishes (IBIDI) (glass bottom, for acetone-fixation) using poly-L-lysine (20 μg/ml, 1 h incubation at 37 °C, SIGMA), labelled with CellTracker™ Blue (7-amino-4-chloromethylcoumarin) (CMAC, 10 μM, 45 min incubation at 37 °C, ThermoFisher) and, after CMAC washing, are pulsed with 1 µg/ml Staphylococcal enterotoxin E (SEE, 45 min incubation at 37 °C, Toxin Technology, Inc). After careful aspiration of the culture medium containing SEE, Jurkat cells are directly added to the microwells, so that IS are formed. CMAC labelling allows to discriminate Jurkat/Raji conjugates. Please refer to the following references for further details, since this IS model has been exhaustively described (Montoya et al., 2002) (Alonso et al., 2011) (Bello-Gamboa et al., 2019). After 30 min-1 h of cell conjugate formation, end-point fixation is performed with chilled acetone for 5 min (*see* **Note 1**).
4. Perform immunofluorescence following standard protocols. Blocking and permeabilization solutions for sequential antibody/phalloidin incubations and washing steps (3x) after each sequential incubation contain saponin (0,1%). The centrosome is labelled with mouse monoclonal anti-γ-tubulin (1/2000 dilution in permeabilization solution, 45 min, room temperature-RT-, Clone GTU-88, SIGMA) and an appropiate secondary antibody coupled to AF546 (1/200 dilution in permeabilization solution, 30 min, RT, Thermofisher). F-actin is labelled with phalloidin AF647 (1/100 dilution in permeabilization solution, 30 min, RT, Thermofisher). CMAC fluorescence does not overlap with these fluorochromes.
5. Perform image capture with confocal microscope. Around 50 optical sections are acquired using a 0.2-0.3 µm Z-step size to have an appropriate axial resolution. Confocal settings we have commonly used are (Leica SP8 confocal microscope): Objective 100x_oil 1.4 NA Zoom_4 Scan Velocity_700Hz_Bidirectional 1024×1024 Pixel Size_0.028 μm Image Size_29.06 μm × 29.06 μm Pinhole_0.7 Optical section_0.704 μm ZStepSize_0.3 μm Z-stack_variable (*top y bottom*) 8 bit Frame Average-2 Line Average-1

## 3. Imaging the IS

### 3.1 Measurement of centrosome polarization index (PI)

1. Open the confocal file in the public, multiplatform software ImageJ or FIJI (https://imagej.nih.gov/ij/) using the “Bio-Formats” import plugin (*see* **Notes 2** and **3**). Save separately the stacks of the different channels in TIF format.
2. Draw the regions of interest (ROI) as seen in Fig. 1 as previously described (Obino et al., 2017) (Bello-Gamboa et al., 2020). Saving all the ROIs by using the “ROI Manager” Image J submenu will allow to recover the ROIs when required (*see* **Note 4**).

**Figure 1.**
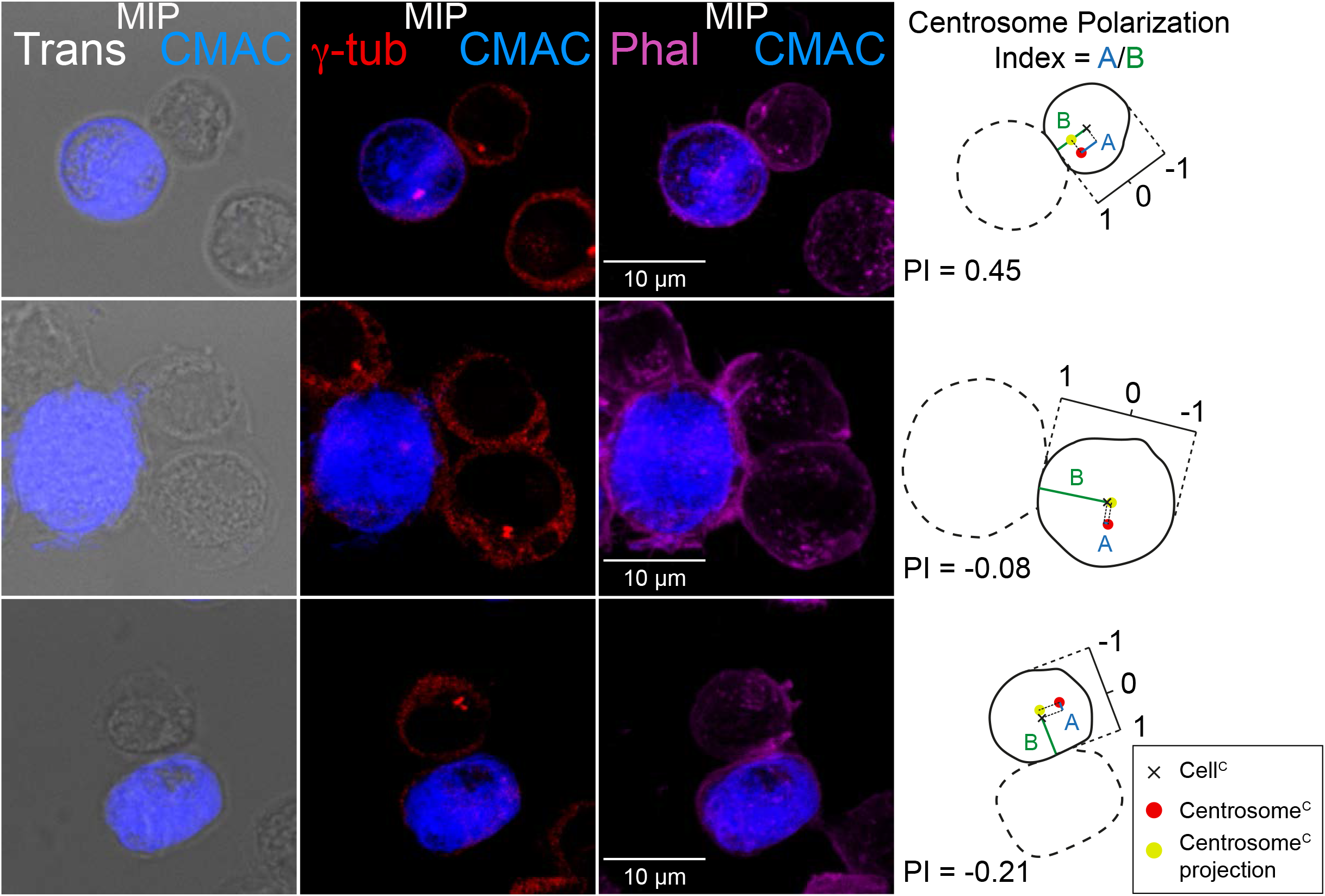
Measurement of centrosome polarization index. Jurkat cells were challenged with CMAC-labelled (blue), SEE-pulsed Raji cells for 1 h, fixed, stained with anti-γ-tubulin AF546 to label the centrosome (red) and phalloidin AF647 to label F-actin (magenta) and imaged by confocal fluorescence microscopy. In the left panels, Maximal Intensity Projection (MIP) of the indicated merged channels (transmittance-Trans-+CMAC, γ-tubulin+CMAC and phalloidin+CMAC) are shown. First row shows an example of a centrosome polarized towards the IS, and second row displays a cell exhibiting a non-polarized centrosome while the lower images show a centrosome in the opposite direction of the IS. The diagrams to the right show the ROIs and the measurements needed for centrosome PI determination: the black crosses label cell^C^; red dots indicate centrosome^C^; yellow dots represent intersections between the B and the projection segments; the segments represent the distances (“A”, blue; “B”, green) used for centrosome PI calculation (A/B). Note that if the intersection between the B and the projection segments is between the cell^C^ and the IS, the value of distance “A” will be positive; otherwise, it must be considered negative. Thus, centrosome PI will vary between +1 (completely polarized) and -1 (completely anti-polarized). Raji and Jurkat cells are labelled with discontinuous and continuous white lines, respectively.
  2.1 Cell shape and cell center of mass.
    - Make the maximal intensity projection (MIP) with the “Z” stacks of the appropriate channel (i.e. phalloidin channel) where cell shape is properly visualized thanks to the cortical F-actin labelling, by selecting “Image” > “Stacks” > “Z Project” > “Max Intensity”. Draw the cell outline (“freehand” selection in ImageJ toolbar) in the MIP (*see* **Note 5**). Save the cell outline as a ROI (cell ROI).
    - Press “Measure” in order to obtain the value of the cell center of mass (cell^C^) (XM and YM coordinates) (*see* **Note 6**).
    - Draw a small rectangle (“rectangular” selection in ImageJ toolbar) inside the cell. Press “Edit” > “Selection” > “Specify” and introduce the values obtained from the cell^C^ in the X and Y coordinates. Select “Scaled units (microns)”. Save it as a ROI (cell^C^ ROI).
  2.2 Analysis of “B” distance.
    - Make a MIP by selecting “Image” > “Stacks” > “Z Project” > “Max Intensity” with the “Z” stacks of the γ-tubulin channel where the centrosome is visualized and establish the centrosome center of mass (centrosome^C^).
    - On the resulting MIP, draw the B segment as the shortest connection between cell^C^ and the IS (“straight line” selection in ImageJ toolbar). Save it as a ROI (B ROI).
    - Press “Measure” to determine “B” segment length (“B distance”).
  2.3 Analysis of “A” value.
    - Draw the projection segment (“straight line” selection in ImageJ toolbar), from the centrosome^C^ perpendicular to the axis defined by the B segment. Save it as a ROI (projection segment ROI).
    - Draw the A segment (“straight line” selection in ImageJ toolbar) from the cell^C^ to the intersection between the B and the projection segments. Save it as a ROI (A ROI).
    - Press “Measure” to determine “A” length. If the centrosome^C^ projection is located between the cell^C^ and the IS, this value will stay positive. However, it must be considered negative if the centrosome^C^ projection is on the opposite side of the cell.
3. Calculate the centrosome PI by dividing “A” distance by “B” distance. The result will vary between +1 (completely polarized) and -1 (completely anti-polarized), thus it will be normalized by the distance between the cell^C^ and the IS (Fig. 1).

### 3.2 Quantification of F-actin at the centrosomal area

1. Define the position of the centrosome and the cellular optical sections to be studied.
  1.1. Open the confocal image capture file in the confocal visualization program (*see* **Note 7**).
  1.2. Determine the “Z” substack in which the centrosome is visually displayed. The appropriate volume to quantify the amount of F-actin around the centrosome is a 2 µm height cylinder with a 2 µm diameter circular base, as previously described (Ibanez-Vega et al., 2019) (Bello-Gamboa et al., 2020). Therefore, adjust the “Z” substack selection to this 2 µm height. After manual selection of the Z optical section containing the maximal signal corresponding to the centrosome (γ-tubulin signal), a 2 µm-high substack of the F-actin channel centered at the centrosome^C^ is defined. Write down the initial and final optical slice. This “centrosomal substack” will comprise 10 optical sections (Z step size= 0.2 µm).
  1.3. Determine the “Z” substack in which all the cell is visually displayed, using phalloidin staining as a reference (*see* **Notes 8** and **9**). Annotate the initial and final optical slice. This will be considered as the “cell substack”.
2. Quantification of centrosomal area F-actin Mean Fluorescence Intensity (MFI) (“C”).
  2.1. Import the confocal image capture file to ImageJ by using the “Bio-Formats” plugin (*see* **Notes 2** and **3**) and visualize the 2 µm-high centrosomal substack of the F-actin channel, as defined in step 1.2.
  2.2. Draw on the file a 2 µm-diameter oval ROI (using “oval” selection in ImageJ toolbar). Then, select “Edit” > “Selection” > “Specify” and choose “Scaled units (microns)”. Adjust ROI length and width to 2 µm and place the circular ROI so its center overlaps the centrosome^C^ (*see* **Note 10**) (Fig. 2). Save it as the centrosomal area ROI (*see* **Note 4**).

**Figure 2.**
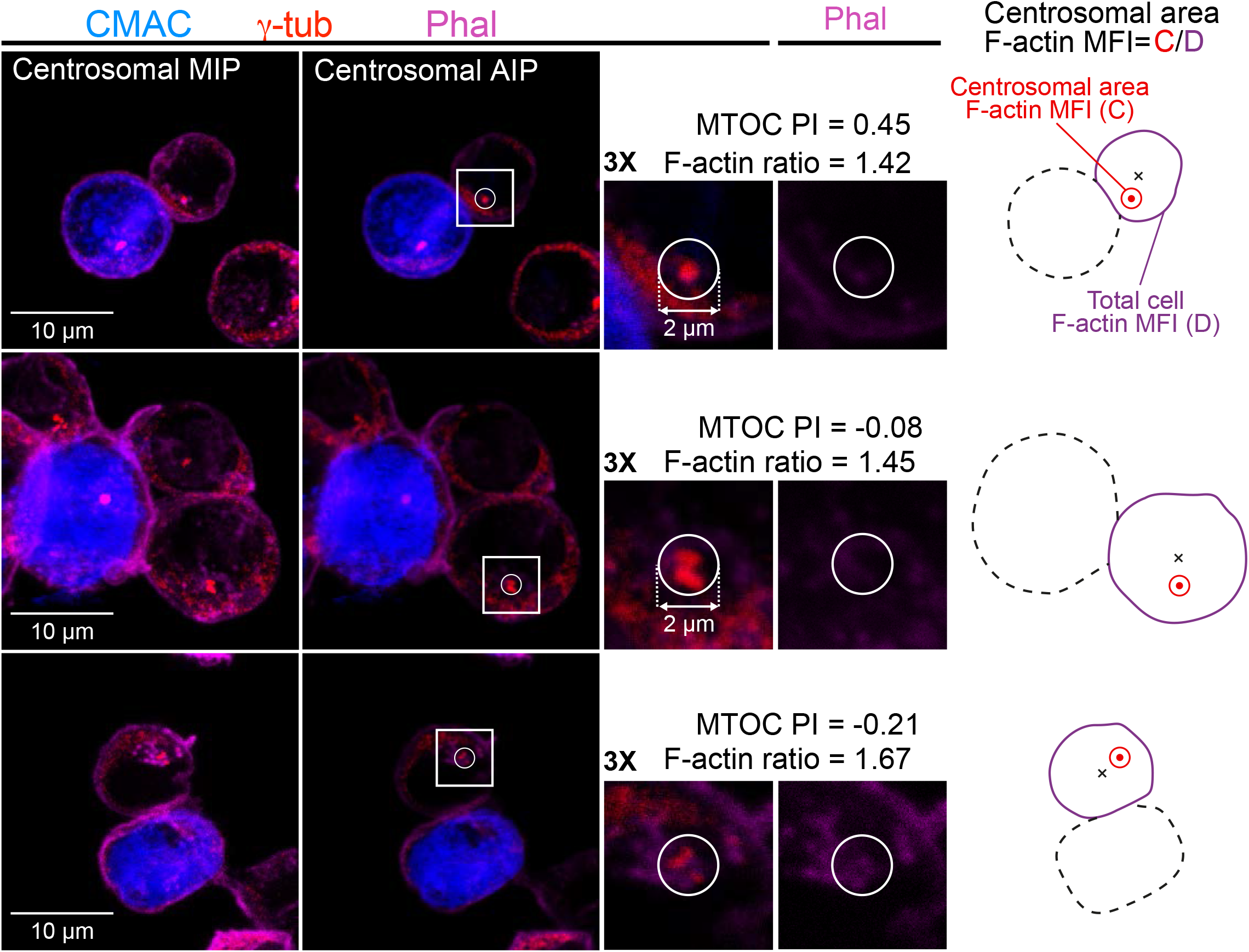
Quantification of centrosomal area F-actin. Jurkat cells were challenged with CMAC-labelled (blue), SEE-pulsed Raji cells for 1 h, then they were fixed, stained with anti-γ-tubulin AF546 to label the centrosome (red) and phalloidin AF647 to label F-actin (magenta) and imaged by confocal fluorescence microscopy. In the left panels, Maximal and Average Intensity Projections (MIP and AIP, respectively) corresponding to the 2 μ m-high, centrosomal substack are shown of the indicated merged channels (CMAC + γ-tubulin + phalloidin) of three representative cells: upper images show a centrosome polarized towards the IS, middle and lower images show centrosomes in the opposite direction of the IS. In the middle panels, 3x enlargements of the centrosomal areas for the three merged channels (left) and for the phalloidin channel (right), defined by white squares in the AIPs, are shown. The 2 µm-diameter ROIs used to calculate centrosomal area F-actin MFI are indicated as white circles. The diagrams to the right indicate the ROIs used in the quantification of the MFI ratio: black crosses, cell^C^; red dots, centrosome^C^; 2 µm-diameter red circles where centrosomal area F-actin MFI (“C”) is measured; purple cell outline where total cell F-actin MFI (“D”) is calculated. Centrosomal area F-actin MFI ratio is calculated as C/D, and the corresponding centrosome PI values are shown. Raji and Jurkat cells are drawn using discontinuous and continuous lines, respectively.
  2.3. Make an Average Intensity Projection (AIP) of the centrosomal substack of the F-actin channel: “Image” > “Stacks” > “Z Project” > “Average Intensity”, by selecting the optical slices defined in step 1.2. Thus, the 2 µm-high “Z” centrosomal substack established in step 1.2 will be delimited. The F-actin AIP of the centrosomal substack is generated (centrosomal F-actin AIP).
  2.4. Select the centrosomal area ROI centered at the centrosome^C^ from step 2.2 in the centrosomal F-actin AIP from step 2.3, and delete the outside of the ROI by selecting: “Edit” > “Clear Outside”.
  2.5. Perform an automatic threshold: “Image” > “Adjust” > “Threshold” (*see* **Note 11**).
  2.6. In order to measure the MFI, select “Analyze” > “Measure” (*see* **Note 12**). The thresholded, centrosomal area F-actin MFI in the centrosomal area ROI is calculated (F-actin MFI in ROI “C”, Fig. 2).
3. Quantification of total cell F-actin MFI (“D”).
  3.1. Make a MIP of the F-actin channel: “Image” > “Stacks” > “Z Project” > “Max Intensity”, by selecting the optical sections defined in step 1.3 (see **Note 13**).
  3.2. Draw the outline of the cell on this MIP (“freehand” selection in ImageJ toolbar) and save it as the cell ROI (*see* **Notes 4, 5, 14** and **15**) (Fig. 2).
  3.3. Make an AIP of the cell substack of the F-actin channel: “Image” > “Stacks” > “Z Project” > “Average Intensity” by selecting the optical sections defined in step 1.3 (see **Note 13**).
  3.4. Select the cell ROI from step 3.2 on this AIP and delete the outside of the ROI by selecting “Edit” > “Clear Outside”.
  3.5. Perform an automatic threshold: “Image” > “Adjust” > “Threshold” (*see* **Note 11**).
  3.6. In order to measure the MFI, select “Analyze” > “Measure” (*see* **Note 16**). The thresholded, total cell F-actin MFI in the cell ROI is calculated (F-actin MFI in ROI “D”, Fig. 2).
4. Determine the centrosomal area F-actin MFI ratio by dividing “C” by “D” (Fig. 2). Thus, centrosomal F-actin measurements will be normalized among samples from different days and different F-actin intensities.

## 4. Discussion, future perspectives and concluding remarks

The method described above is not automated and, therefore, to perform the quantification the user must select single cells forming IS one by one, which may make it difficult to evaluate results in a fully unbiased form and also may be time-consuming to perform for an untrained user. However, a trained user may analyze up to 20-30 synapses per hour. To prevent biases, please try to randomly select as many synaptic conjugates from different fields as possible. In addition, we have tried artificial intelligence programs such as the NisAR supplement Ai (NIKON) to automatically identify and select synaptic conjugates for subsequent analysis. Unfortunately, in our experience this approach cannot distinguish canonic IS from irrelevant cell contacts or cell aggregates randomly produced by the high cell concentrations used to favour IS formation (Bello-Gamboa et al., 2019). Improvement of the artificial intelligence algorithms will contribute to solve this problem. Evaluation of the cup-shape profile of the effector T lymphocyte and/or lamellipodium formation observed in the transmittance channel may help to identify IS. A criterion we recommend to unambiguously select canonic IS and distinguish them from irrelevant cell contacts is to use the F-actin probe phalloidin (Fig. 1 and Fig. 2), since it is known that F-actin accumulates at the lamellipodium in the synaptic contact area (please refer to Introduction). CMAC labelling facilitates identification of Jurkat/Raji conjugates.

In order to somewhat automate image analyses and save time we have used the ImageJ “Macro” language (IJM), that is a scripting language built into ImageJ that allows controlling many aspects of this software. A macro is a simple program that automates a series of ImageJ commands and may facilitate dealing with image stacks. Programs written in the IJM, or macros, can be used to perform sequences of actions in a fashion expressed by the program’s design. The easiest way to create a macro in ImageJ is to record a series of commands using the command recorder (“Plugins” > “Macros” > “Record”). A macro is saved as a text file and executed by selecting a menu command, by pressing a key or by clicking on an icon in the ImageJ toolbar. Please refer to the excellent ImageJ tutorials https://imagej.nih.gov/ij/developer/macro/macros.html. We have found the following macros in txt format very useful:

run(“Make Substack…”,”slices=X-Y”);

run(“Z Project…”, “projection=[Average Intensity]”);

The first macro simplifies substack generation (changing X and Y characters by the optical section number) and the second macro directly generates the AIP from the former substack or a different substack in a single operation. Thus, these macros could be used in steps 2.3 and 3.3, and can be used separately or together, as required.

Several authors, including ourselves, operationally used the AIP of a 2 μm-high, F-actin confocal substack centered at the centrosome on an arbitrarily-defined, 2 µm-diameter circular region of interest (ROI) centered at the centrosome, to measure centrosomal area F-actin reorganization triggered by Th (Bello-Gamboa et al., 2019) and B lymphocytes (Obino et al., 2016) (Ibanez-Vega et al., 2019) IS formation. However, it cannot be excluded that other organelles, such as the Golgi (Colon-Franco et al., 2011) or endosomes/MVB (Calabia-Linares et al., 2011) (Ueda et al., 2011), that are competent in reorganizing F-actin or tubulin cytoskeleton, may be included in this area, and thus we cannot exclude the contribution of Golgi/MVB to the F-actin reorganization at the centrosomal area during centrosome polarization. The imaging techniques used in these reports (confocal microscopy, total internal reflection microscopy -TIRFM-, epifluorescence microscopy plus deconvolution) do not provide enough resolution to discriminate between F-actin assembly at the pericentrosomal matrix or other membrane-bound organelles, that may be included in the 2 µm-diameter centrosomal area ROI (Obino et al., 2016) (Ibanez-Vega et al., 2019) (Bello-Gamboa et al., 2019). Indeed, in the future, emerging and promising imaging techniques applied to living cells, harboring high spatio-temporal resolution such as lattice light-sheet microscopy (LLSM) (Ritter et al., 2015) (Fritzsche et al., 2017), combined with non-diffraction limited, super-resolution microscopy (Fritzsche et al., 2017) (Calvo and Izquierdo, 2018), may contribute to a better definition of centrosomal area F-actin structure and function, as it occurred for the four distinct synaptic F-actin networks in the canonical IS (Hammer et al., 2018)

(Blumenthal and Burkhardt, 2020). By using these approaches, it will be interesting to analyse, for instance, whether centrosomal F-actin clearing that occurs during centrosome polarization to the IS is a reversible event, as it occurs with cortical F-actin in IS made by CTL (Ritter et al., 2015) (Ritter et al., 2017). In the future, this analysis will allow extending the knowledge regarding the contribution of centrosomal area F-actin to polarized secretion obtained in B and Th lymphocytes IS (Obino et al., 2016) (Bello-Gamboa et al.) to CTL and NK cells IS, due to the similarities existing among all these IS. In addition, all these approaches together with our analysis will enable the study of the contribution of centrosomal area F-actin to polarized secretion during directional and invasive cell migration, both of lymphoid and non-lymphoid cells (Calvo and Izquierdo, 2021). This would clarify whether centrosomal area F-actin reorganization occurs only during IS formation or contributes also to other biologically relevant cellular polarization processes. This analysis will enable to address how the distinct cortical and centrosomal F-actin networks are regulated, how these networks integrate into cell surface receptor-evoked signalling networks, as well as their interconnections with tubulin cytoskeleton, that constitute an intriguing and challenging biological issue (Dogterom and Koenderink, 2019) (Calvo and Izquierdo, 2021).

## 5. Notes

1. Acetone precipitates FCS protein, thus a previous wash in warm culture medium without FCS prior to fixation is recommended. Acetone (and acetone vapour) dissolves plastic, thus keep the microwell with fixative on ice at 4° C, and the microwell plastic lid should be removed. Recommended maximal fixation time is 5 min, since longer times dehydrate samples in excess (producing cell shrinking and cell shape changes). Acetone fixation does not allow so clean staining of centrosome with anti-γ-tubulin as methanol fixation does, although centrosome staining still is evident (Figs. 1 and 2). We noticed this occurs regardless of blocking and permabilization conditions and the number of washings steps during immunofluorescence. Paraformaldehyde or glutaraldehyde does not allow staining of centrosome with anti-γ-tubulin. An important issue is that F-actin staining with phalloidin is not compatible with methanol fixing, whereas it is compatible with acetone fixation. Thus, we have chosen acetone fixation since it constitutes a good compromise to circumvent these caveats (Abrahamsen et al., 2018). As an alternative, some authors have used glutaraldehyde fixation instead, which enables proper F-actin labelling, but not anti-γ-tubulin staining (Obino et al., 2016) (Ibanez-Vega et al., 2019). In this condition, centrosome position was indirectly inferred with anti-α-tubulin staining, as the brightest point where the microtubules converge by using an appropriate threshold. We believe our approach is more direct indeed.
2. If a channel is not properly visualized, you can modify its optical parameters in “Image” > “Adjust” > “Brightness/Contrast”. Please do not apply these changes especially during F-actin measurement, otherwise intensity values will be modified.
3. In order to improve and synchronize the visualization of the different channels you can select “Window” > “Tile” and “Analyze” > “Tools” > “Synchronize windows”. Also, you can visualize and merge the channels you are interested in by selecting “Image” > “Color” > “Merge Channels”.
4. To open the ROI manager press “Analyze” > “Tools” > “ROI manager”. Select “Show All” so afterwards you can always recover and visualize the previously saved ROIs. Leave this window open for next steps.
5. In ImageJ the drawing is performed with the “freehand” tool using a proper channel to outline the cell (i.e. phalloidin). Other analysis software includes an “autodetect” option. For the study of the IS this option is useful only if the effector cell being studied has a clear differential staining (e.g. expresses a reporter gene such as GFP).
6. The first time the analysis is performed, it is necessary to establish the parameters that are going to be measured: “Analyze” > “Set Measurements” and select “Center of mass”.
7. Always use the same computer screen in order to avoid discrepancies in ROI drawing and Z substack selection and minimize the error between measurements.
8. If cell shape differs considerably from one “Z” slice to another, divide the “Z” slices into different substack groups.
9. In order to avoid discrepancies and minimize the error between measurements, do not select “Z” stacks where the F-actin is barely seen.
10. The circular ROI must not include the IS F-actin, which may happen in cells with a highly polarized centrosome towards the IS. Then the centrosomal area may overlie the synaptic membrane rich in cortical F-actin and bias the measurements.
11. Use “Default” setup and do not select “Apply”, since it would affect the MFI measurements.
12. The first time the analysis is performed, it is necessary to establish the parameters that are going to be measured: “Analyze” > “Set Measurements” and select “Limit to threshold”, “Area” and “Mean gray value”.
13. If different substack groups were selected in step 1.3, make a MIP for each substack.
14. Be careful not to include F-actin from nearby cells.
15. Draw a ROI for each MIP of step 3.1.
16. If different substacks were selected in 1.3, MFI value must be computed by calculating a weighted average of the values obtained by following steps 3.2-3.6 in the different substack groups and considering the number of “Z” slices in each substack group.

## ABBREVIATIONS

AIP: average intensity projection
APC: antigen-presenting cell
BCR: B-cell receptor for antigen
^C^: center of mass
CMAC: CellTracker™ Blue (7-amino-4-chloromethylcoumarin)
cSMAC: central supramolecular activation cluster
CTL: cytotoxic T lymphocytes
Dia1: diaphanous-1
dSMAC: distal supramolecular activation cluster
F-actin: filamentous actin
(FCS): fetal calf serum
FMNL1: formin-like 1
GFP: green fluorescent protein
IS: immunological synapse
LLSM: lattice light-sheet microscopy
MFI: mean fluorescence intensity
MHC: major histocompatibility complex
MIP: maximal intensity projection
MTOC: microtubule-organizing center
MVB: multivesicular bodies
NK: natural killer
PI: polarization index
PKCδ: protein kinase C δ isoform
pSMAC: peripheral supramolecular activation cluster
ROI: region of interest
RT: room temperature
SEE: Staphylococcal enterotoxin E
SMAC: supramolecular activation cluster
TCR: T-cell receptor for antigen
Th: T-helper
TIRFM: total internal reflection microscopy
TRANS: transmittance

## Author Contributions

S.F., I.S., and M.B. wrote the manuscript and prepared the figures. V.C. and M.I. conceived the manuscript and the writing of the manuscript and approved its final content. Conceptualization, V.C. and M.I; writing original draft preparation, M.I.; reviewing and editing, V.C. and M.I.

## Acknowledgements

This research was funded by grants from the Spanish Ministerio de Ciencia e Innovación, (PID2020-114148RB-I00) to MI, which was in part granted with FEDER funding (EC), corresponding to Programa Estatal de Generación de Conocimiento y Fortalecimiento Científico y Tecnológico del Sistema de i+d+i y de i+d+i Orientada a los Retos de la Sociedad.

